# Protein expression profiles in brain organoids are more similar to those in human brain parenchyma than in mouse brain parenchyma

**DOI:** 10.1101/2023.09.12.557448

**Authors:** Tyler J. Wenzel, Darrell D. Mousseau

## Abstract

Human brain organoids are emerging as relevant models for the study of human brain health and disease. However, it has not been shown whether human brain organoids exhibit a proteoform profile similar to the human brain. Herein, we demonstrate that unguided brain organoids exhibit minimal batch-to-batch variability in cell composition and metabolism when generated from induced pluripotent stem cells (iPSCs) derived from male-female siblings. We then show that profiles of select proteins in these brain organoids are more similar to autopsied human cortical and cerebellar profiles than to those in mouse cortical samples. Brain organoids derived from sibling iPSCs do not exhibit any sex differences in protein proportions. By benchmarking human brain organoid proteoforms against human parenchymal tissue, we establish the foundation for future studies that could investigate, for example, how well brain organoids can model any of the known sex-dependent differences in cellular function, including responses of drug-receptor interactions.

**Highlights:**

- Brain organoids (BOs) display protein banding similar to human parenchymal lysates
- Protein banding differs between mouse and human brain parenchyma lysates
- Sibling-derived BOs have similar cell composition and metabolism at day 90
- Sibling-derived BOs exhibit similar protein banding at day 90

## 1 Introduction

While human brain organoids (BOs) were first described over a decade ago (Eiraku et al., 2008), there is still a limited understanding about which features of the human brain these cultures mimic, and whether they can replace the use of animals to provide insight into health and disease (Wenzel et al., 2023). Benchmarking human BOs, where ethically possible, to human brain tissue is critical because interspecies differences complicate the interpretation of data (Bellinger et al., 2011; Jensen and Little, 2023; Maksour and Ooi, 2023; Webster et al., 1995). Furthermore, the quality control of antibodies by suppliers often relies on recombinant proteins or immortalized cell lines as positive controls for antibodies. As suppliers using recombinant proteins and cell lines do not take into account post-translational modifications during validation, the protein banding pattern may not be similar to complex tissues such as brain parenchymal tissues and BOs, complicating interpretation of immunoblotting data (Haytural et al., 2019; Rosell et al., 2020). Key evidence to validate BOs as a model would be to demonstrate that BOs exhibit protein banding patterns similar to human parenchymal tissue.

BOs are not often used for immunoblotting experiments (Zhao et al., 2022). In part, this may be because stringent protocols without the use of concentrated proteinaceous capsules are required for replicable results (Hernández et al., 2022; Wenzel et al., 2023). As such, very few studies have compared human parenchymal tissue and BOs using immunoblotting analysis. We only identify two studies that have compared these human tissues with the mouse parenchyma (Lancaster et al., 2013; Sriram et al., 2020), and these studies only investigated a few proteins of interest. A wider comparison of cell and synapse markers between primary parenchymal tissue and BOs would be beneficial for future studies (Kroon et al., 2022).

Herein, we compare the profiles of select proteins in human BOs, human cortical (Ctx) and cerebellar (Cb) autopsy brain tissues and mouse Ctx tissues. We demonstrate that banding pattern of several human BOs proteins are more similar to human Ctx or Cb tissues than mouse Ctx tissues, indicating that the proteins in human BOs are processed in a manner similar to the proteins in the human parenchyma. Furthermore, we compare the levels of several proteins in human BOs, human and mouse Ctx tissue as well as human Cb tissue derived from male and female donors. BOs were generated from the induced pluripotent stem cells (iPSCs) of female-male sibling donors because familial cell lines are known to result in more similar organoids (Jourdon et al., 2023). Our data corroborate these previously reported data as we observed limited variability between our batches of human BOs, and interestingly, these *in vitro* cultures display less protein variability between samples than the human and mouse parenchymal tissues. Human BOs exhibit human brain-like properties, and our study adds to the evidence that these stem cell-based cultures may help bridge the gap between preclinical and clinical studies.

## 2 Materials and methods

### 2.1 Resource availability

#### Materials availability

This study used reagents from commercial sources, and thus did not generate any new reagents. Requests for materials are promptly reviewed by the University of Saskatchewan to determine if the request is subject to any intellectual property, confidentiality obligations, or material transfer agreements. Data and materials may be subject to material transfer agreements.

#### Data and code availability

All raw data are available upon request. To facilitate data sharing, we include images of full immunoblot membranes in this manuscript. No code was used for data analysis.

### 2.2 Human and mouse tissues

The use of human autopsy tissues in this study were obtained from the Douglas-Bell Canada Brain Bank (McGill University, Canada) and covered by the University of Saskatchewan’s Research Ethics Office Certificate of Approval ‘Bio 06-124’ (principal investigator: Darrell D. Mousseau).

Animal experiments were approved by the University of Saskatchewan’s Animal Research Ethics Board (Protocol No. 20060070) and was performed in accordance with the Canadian Council on Animal Care standards. C56Bl/6J mice (Strain No. 000664) were received from the Jackson Laboratory (Bar Harbor, Maine, USA), maintained in the animal facility, housed in individually ventilated cages, and handled on a clean bench in a specific pathogen-free environment. All mice had free access to rodent feed and reverse osmosis water in a temperature-controlled room (21-22°C) with reverse lighting cycle (12 h dark/light). Mice were euthanized at the age described in **Table 2**, their brain tissues collected and the cortical region was isolated for immunoblotting.

### 2.3 Antibodies and reagents

L-ascorbic acid (LAA), ethylenediaminetetraacetic acid, heparin, laminin from Engelbreth-Holm-Swarm murine sarcoma, Lowry assay kit (Peterson’s modification), poly-D-lysine hydrobromide, protease inhibitor cocktail (Cat#P8340), sodium selenite, and Terg-A-Zyme detergent were purchased from Millipore Sigma (Oakville, ON, Canada). Transforming growth factor-β1 (TGF-β1; cat #100-21) was purchased from Peprotech (Cranbury, NJ, USA). Recombinant human transferrin (cat#777TRF029) was purchased from InVitria (Fort Collins, CO, USA). Radioimmunoprecipitation assay buffer (RIPA) 10x and the ROCK inhibitor Y-27632 (cat# 13624S) were purchased from Cell Signaling Technologies (Whitby, ON, Canada). BrainPhys™ Neuronal Medium was obtained from STEMCELL Technologies (Vancouver, BC, Canada). A full list of antibodies and their suppliers is provided in **Supplementary Table 1**. All other reagents were sourced from Fisher Scientific (Ottawa, ON, Canada).

### 2.4 Inducible pluripotent stem cell (iPSC) maintenance

UCSD086i-6-3 (86i, male) and UCSD087i-6-4 (87i, female) iPSC lines, which are derived from a male-female sibling pair, were purchased from WiCell (Madison, WI, USA) and confirmed to be karyoptypically normal by the provider. These were used in all experiments and were cultured in feeder-free conditions on 6-well tissue culture plates coated with Matrigel™ hESC-qualified matrix (Corning™ 354277) or Geltrex™ hESC-qualified matrix and maintained at 37 °C in humidified 5% CO_2_ and 95% air atmosphere.

### 2.5 Generation of unguided brain organoids

Human unguided BOs were generated as described elsewhere (Wenzel, 2022). Briefly, iPSCs were incubated with 0.5 mM ethylenediaminetetraacetic acid (EDTA) for five minutes. They were then removed from the plate with a cell scraper, centrifuged, and resuspended at 9 x 10^4^ iPSCs/ml in embryoid body (EB) maintenance media with 10 μM Y-27632 inhibitor. ∼9000 iPSCs were seeded in a 96-well ultra-low attachment round-bottom plate. 48 hours later (day 2), 100 μl of EB maintenance medium were added to each well. On day 5, using a 200 μl wide-bore pipette tip and ensuring minimal transfer of EB maintenance medium, individual EBs were transferred to separate wells of a 24-well ultra-low attachment plate containing 500 μl of induction media. On day 7, media was replaced with 500 μl of ice-cold expansion media and on day 10, 1 ml of maturation media was added to each well and incubated until day 17 at 37 °C in humidified 5% CO_2_ and 95% air atmosphere on an orbital plate shaker set at 0.118 *g*. Once a week, media was replaced with 750 μl of maturation media, followed by an additional 500 μl being added four days later. BOs in culture were harvested between day *in vitro* 10 and 90 (as indicated on figures) and then protein extracts were assessed for protein content and prepared for standard denaturing SDS-PAGE immunoblotting. Brightfield images of BOs were taken with an Olympus CK53 microscope.

### 2.6 Immunoblotting

Five BOs were pooled and homogenized in RIPA buffer containing protease inhibitor cocktail. Samples were triturated with a one ml pipette, homogenized using a 22-gauge needle, and sonicated. Protein concentration was quantified using the Lowry (Folin-Ciocalteu phenol reagent) assay and equalized to 0.5 μg/μl in 1% loading buffer (0.2 M Tris pH 6.8, 40% glycerol, 8% sodium dodecyl sulfate, 20% β-mercaptoethanol, 0.4% bromophenol blue). Samples were not heated to prevent protein aggregation. Resolved proteins (10 or 15 µg per lane, as indicated in **Table 1**) were electroblotted onto a nitrocellulose membrane and blocked in 5% bovine serum albumin (BSA) in TRIS-buffered saline (TBS: 250 mM Tris pH 7.4, 1.37 M NaCl) for one hour. Membranes were the washed and incubated overnight at 4 °C with primary antibodies (**Table 1**) diluted in 5% BSA in TBS-T (TBS with 0.1% Tween®20). After three washes with TBS-T over 30 min, secondary fluorescently-labelled antibodies (**Table 1**) were added for one hour, followed by three washes. Proteins were visualized with a LI-COR Odyssey® Imager and analyzed with manufacturer’s software (Image Studio 5.3.5, LI-COR Biosciences, Lincoln, NE, USA).

**Table 1:** List of antibodies and reagents used.

| Target | Catalogue Number | Blotting Dilution | Amount of protein loaded |
| --- | --- | --- | --- |
| **Primary Antibodies** |  |  |  |
| Rabbit anti-β-actin | Cell Signaling Technology<br>Cat#4970 | 1:5,000 | Fig. 1: 15 μg<br>Fig. 5: 10 μg |
| Rabbit anti-GAPDH | Cell Signaling Technology<br>Cat#5174 | 1:2000 | Fig. 1: 15 μg<br>Fig. 5: 10 μg |
| mouse anti-GFAP | Millipore Sigma<br>Cat#3893 | 1:1,000 | Fig. 1: 15 μg<br>Fig. 4: 10 μg |
| Rabbit anti-IBA1 | Abcam<br>Cat# ab178846 | 1:1,000 | Fig. 1: 15 μg<br>Fig. 3: 10 μg |
| rabbit anti-OLIG2 | Millipore Sigma<br>Cat#AB9610 | 1:1,000 | Fig. 1: 15 μg<br>Fig. 4: 15 μg |
| Rabbit anti-P2RY12 | Atlas Antibodies<br>Cat#HPA014518 | 1:1,000 | Fig. 3: 20 μg |
| Mouse anti-NeuN | Millipore Sigma<br>Cat#MAB377 | 1:2,000 | Fig. 1: 15 μg<br>Fig. 2: 10 μg |
| Rabbit anti-TLR4 | Santa Cruz Biotechnology<br>Cat#sc-293072 | 1:1,000 | Fig. 5: 10 μg |
| Mouse anti-TMEM119 | BioLegend<br>Cat#A16075D | 1:500 | Fig. 1: 15 μg<br>Fig. 3: 20 μg |
| Rabbit anti-TUBB3 | Millipore Sigma<br>Cat#T2200 | 1:5,000 | Fig. 1: 15 μg<br>Fig. 2: 10 μg |
| Rabbit anti-SYN1 | Cell Signaling<br>Technology<br>Cat#5297 | 1:1000 | Fig. 1: 15 µg<br>Fig. 2: 10 µg |
| Secondary Antibodies |  |  |  |
| IRDye® 680RD anti-rabbit IgG | LI-COR<br>Biosciences<br>Cat#926-68071 | 1:20,000 | N/A |
| IRDye® 680RD anti-mouse IgG | LI-COR<br>Biosciences<br>Cat#926-68070 | 1:20,000 | N/A |
| IRDye® 800CW anti-rabbit IgG | LI-COR<br>Biosciences<br>Cat#926-32211 | 1:20,000 | N/A |
| IRDye® 800CW anti-mouse<br>IgG | LI-COR<br>Biosciences<br>Cat#926-32210 | 1:20,000 | N/A |
| Reagent | Catalogue Number |  |  |
| Growth Factors |  |  |  |
| Thermostable FGF2 | ThermoFisher Scientific Cat#PHG0369 |  |  |
| TGF-β1 | PeproTech Cat#100-21 |  |  |

Abbreviations: βIII-Tubulin (TUBB3), basic fibroblast growth factor (FGF2), glial fibrillary acidic protein (GFAP), glyceraldehyde 3-phosphate dehydrogenase (GAPDH), immunoglobulin G (IgG), ionized calcium-binding adapter molecule 1 (IBA1), neuronal nuclei (NeuN), oligodendrocyte transcription factor 2 (OLIG2), purinergic Receptor P2Y12 (P2RY12), synapsin 1 (SYN1), toll-like receptor 4 (TLR4), transforming growth factor (TGF), transmembrane protein 119 (TMEM119).

**Table 2:** Donor information.

| **Human Cortex (From Neurocognitively Normal Donors)** |  |  |  |
| --- | --- | --- | --- |
| **Donor Sex** | **Donor Age** | **Apolipoprotein E Alleles** | **Cause of Death** |
| Male | 61 years old | ε3/ε3 | Pancreatic metastasis |
|  | 77 years old | ε3/ε3 | Information not available |
|  | 60 years old | ε3/ε3 | No discernable cause |
| Female | 59 years old | ε3/ε3 | Chronic obstructive pulmonary disease |
|  | 75 years old | ε3/ε3 | Pulmonary edema |
|  | 91 years old | ε3/ε3 | Information not available |
| **C57Bl/6J Mouse Cortex** |  |  |  |
| **Donor Sex** | **Donor Age** |  |  |
| Male | Postnatal week 11 |  |  |
|  | Postnatal week 14 |  |  |
|  | Postnatal week 15 |  |  |
| Female | Postnatal week 11 |  |  |
|  | Postnatal week 14 |  |  |
|  | Postnatal week 15 |  |  |

### 2.7 Multielectrode array recording

BOs were plated on a MaxOne high-density multiectrode array (MaxWell Biosystems, Zurich, Switzerland) following manufacturer’s instructions. In brief, electrodes were treated with 1% (w/v) Terg-a-zyme for 2 hours. Next, electrodes were coated with 0.1 mg/ml poly-D-lysine dissolved in a borate buffer for 1 hour, washed three times, and then coated with 0.04 mg/ml laminin dissolved in culture media for 1 hour. BOs were then allowed to attach to the multielectrode array over two weeks in a CO_2_ incubator set to 37 °C. Media was replaced twice a week with 600 μl of BrainPhys™ Neuronal Medium. After a two-week incubation, neural network activity was measured using the Activity Scan Assay followed by the Network Activity Assay in the MaxLab Live software (MaxWell Biosystems).

### 2.8 Cell viability

The PrestoBlue assay was used (following manufacturer’s instructions) to measure the reduction of resazurin to resorufin (a red fluorescent compound) by living cells in organoids. Data are presented as fluorescence signal of resorufin.

### 2.9 Data analysis

GraphPad Prism 9.2 software was used for all statistical analyses. Data (mean ± standard deviation (SD)) were analyzed using the (1) Mann-Whitney test, or the (2) Kruskal-Wallis test, followed by the Dunn’s or the Sidak’s post-hoc test. Significance was established at *p* < 0.05. For mouse and human parenchyma data, each data-point was derived from a different donor (see **Table 2** for donor information). For BO data, the individual data-points were derived from batches of organoids grown on different days.

## 3 Results

### 3.1 Brain organoids reproducibly display the visual characteristics of normal development

We compared the development of unguided BOs generated from the 86i (male) and 87i (female) familial iPSC lines. **Figure 1A** shows BOs generated from the 86i and 87i iPSCs displayed all the expected visual characteristics of normal unguided BO development (Lancaster and Knoblich, 2014). Five BOs were collected every 10 days for 90 days, and the levels of cell and synaptic markers were measured using standard immunoblotting approaches. **Figure 1B** shows that BOs developed cell markers at comparable time-points. At day 90, BOs consistently expressed comparable levels of total protein (**Figure 1C**) as well as housekeeping genes TUBB3, β-actin and GAPDH (**Figure 1D**) regardless of the batch of iPSCs used to generate the BOs. The size of organoids as measured by total protein (**Figure 1G**) as well as their ability to reduce resazurin (**Figure 1H**) were also similar between batches.

**Figure 1:**
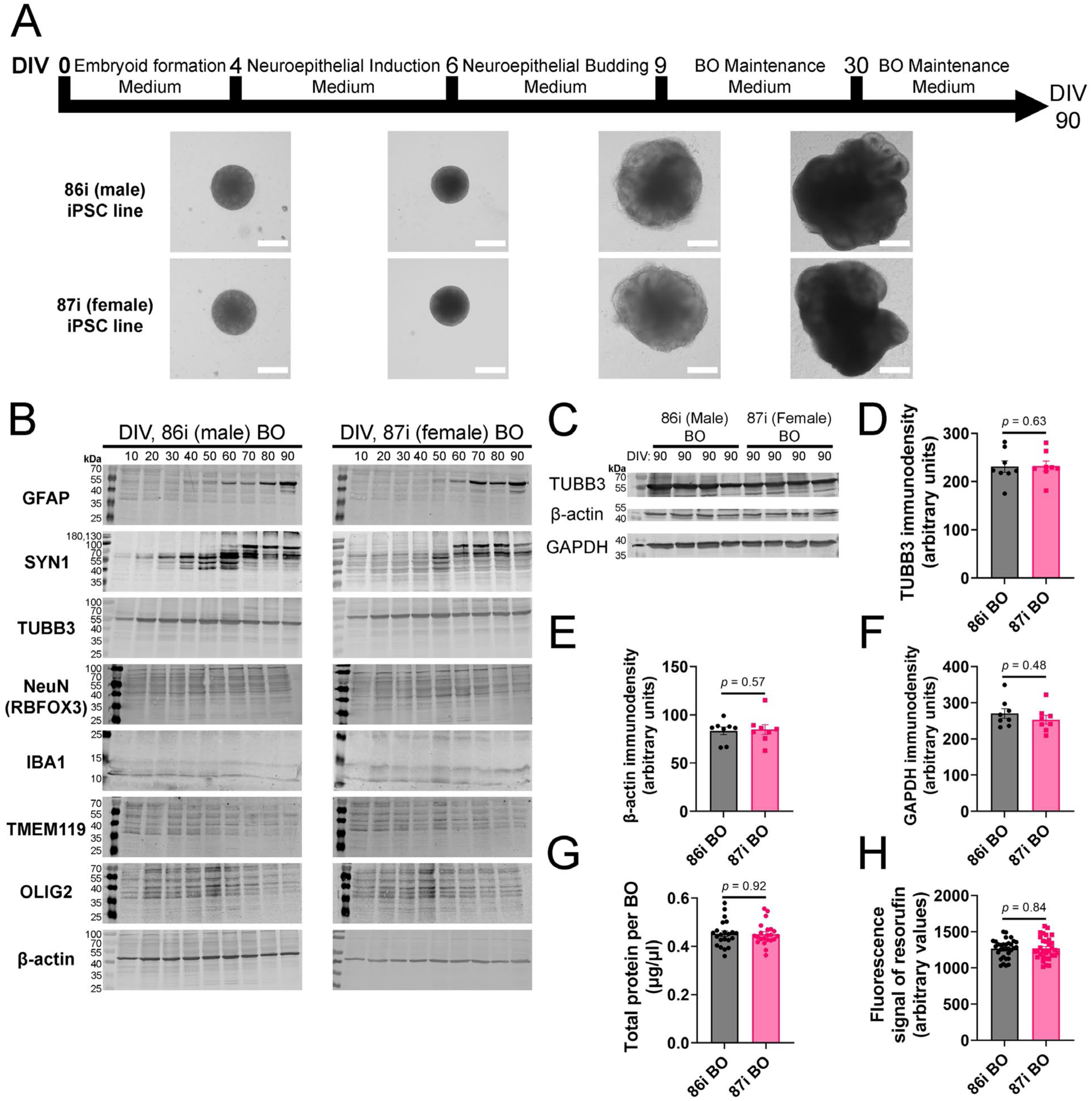
BOs were grown as described in Wenzel (2022), and (A) BOs derived from 86i and 87i cell lines exhibit all visual markers of proper development. (B) Expression of GFAP, SYN1, TUBB3, NeuN, IBA1, TMEM119, OLIG2 and β-actin across 90 days of development were measured. Immunoblotting analysis indicates that BOs become more complex as they are cultured for longer periods of time, as exemplified by the presence and maturation of GFAP and SYN1. At day 90, (C-F) the levels of housekeeping proteins TUBB3, β-actin, and GAPDH are consistently expressed at similar levels, (G) as are the levels of total protein, indicating these cultures grow to similar sizes and complexities at similar rates. (H) The ability of BOs to metabolize resazurin into resorufin is also similar at day 90, indicating similar metabolic activity. (D-H) Data (means ± SD) analyzed according to Mann-Whitney test. (B-F) BO data were derived by pooling five organoids. The five pooled BOs were from the same batch, but each sample (*i.e.,* lane or data-point) was from a different batch of five organoids (batches defined as BOs generated on different days from iPSCs of a different passage number). (G,H) Total protein and resorufin data-points represent individual organoids.

### 3.2 Differences and similarities of protein banding patterns for select proteins in brain organoids as well as in human and mouse brain parenchymal tissues

Limited studies have directly compared the immunoblotting results of human BOs to human and mouse parenchymal tissue. To demonstrate BOs can recapitulate aspects of the human brain, we compared the immunoblotting results from experiments using BOs, human and mouse Ctx tissues and human Cb tissues. As BOs expressed a range of cell markers and synaptic proteins at day 90 (**Figure 1B**), this time-point was selected for the remaining experiments in this study. To demonstrate protein banding similarities between samples, full immunoblots are shown in **Figures 2-5**, and they demonstrate that BOs exhibit protein banding patterns that are more similar to the human Ctx or Cb tissue than the mouse Ctx tissue. Immunoblot images were taken at the same time with the same imaging settings, so the banding intensity is comparable visually.

**Figure 2:**
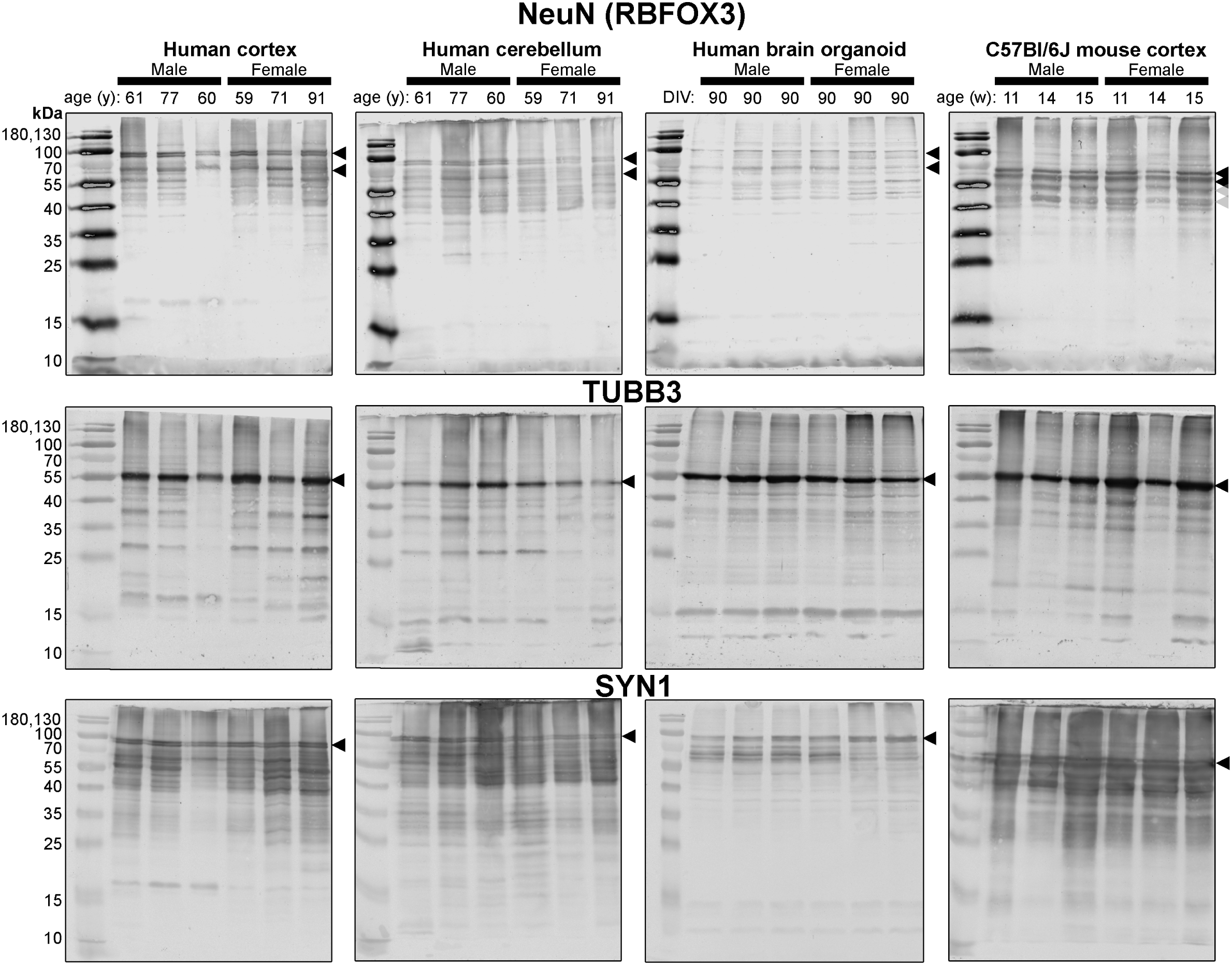
Immunoblots of neuron-associated proteins NeuN (*top*), TUBB3 (*middle*) and SYN1 (*bottom*). Protein homogenate for primary mouse and human tissues were derived from different donors (**Table 2**), while BO homogenate were derived by pooling five organoids. The five BOs were from the same batch, but each sample (*i.e.,* lane) was from a different batch of five organoids (batches defined as BOs generated on different days from iPSCs of a different passage number). Samples of human Ctx and Cb, BOs as well as mouse Ctx were simultaneously ran on an 12% SDS-PAGE gel, transferred onto a nitrocellulose membrane, and treated with antibodies in the same containers. The amount of protein loaded for each sample and the antibodies used are indicated in **Table 1**. Membranes images of the same protein were also taken at the same time as a single image, and only cropped and digitally repositioned for publication. No post processing was done on any image, and so the immunodensity of bands are visually comparable from one blot to the next. We note that TUBB3 was probed overtop of the IBA1 immunoblot shown in Figure 3, but the 55 kDa band was clearly the only new band detected by TUBB3 antibodies, which is consistent with the TUBB3 immunoblot in Figure 1. Arrowheads indicate the specific bands used for estimations of protein expression for Figures 6-7. *Abbreviations: days in vitro (DIV), week (w), year (y), neuronal nuclei (NeuN), β3-tubulin (TUBB3), synapsin 1 (SYN1)*.

First, we immunoblotted for neuron-associated proteins (**Figure 2**). NeuN antibodies detect a 110 kDa protein doublet in human tissues, whereas the doublet resolves between 55-70 kDa in mouse Ctx tissue (**Figure 2**, *top*). NeuN antibodies detect a protein between 55-70 kDa in human tissues, but no doublet is observed. TUBB3 exhibits a similar protein banding pattern between all tissues tested (**Figure 2**, *middle*). The SYN1 protein doublet is detected between 70-100 kDa in human Ctx and Cb tissues as well as BOs (**Figure 2**, *bottom*). In contrast, the SYN1 protein doublet is detected between 55-70 kDa in mouse Ctx tissues.

Next, we probed for proteins associated with the resident immune cells of the brain, namely microglia (**Figure 3**). IBA1 antibodies detected a protein at 17 and 15 kDa in human Ctx tissues, whereas proteins were detected at 12 and 15 kDa in the other tested tissues (**Figure 3**, *top*). TMEM119 antibodies detect a 55 kDa protein in mouse Ctx tissues and a triplet between 40-55 kDa in both human brain and BO tissues (**Figure 3**, *middle*). We also detect a 17 kDa band on the TMEM119 blots of human brain parenchymal tissues. The P2RY12 antibodies detect protein at 17 kDa and around 40 kDa in human tissues (human brain and BOs), whereas only a stronger 40 kDa band is detected in a subset of mouse Ctx samples (**Figure 3**, *bottom*). We then immunoblotted for proteins GFAP and OLIG2 (**Figure 4**), which are commonly associated with the macroglia astrocytes and oligodendrocytes, respectively. GFAP is often described as a 55 kDa protein but has several known isoforms between 35-55 kDa (Kamphuis et al., 2014, 2012). GFAP antibodies detected several isoforms in human brain and BOs, but only one isoform in mouse Ctx tissues (**Figure 4**, *top*). While OLIG2 antibodies detected two bands around 40 kDa in all samples, BO immunoblots displayed more bands at higher molecular weights (**Figure 4**, *bottom*). We note a 17 kDa band that appears in OLIG2 blots of human parenchymal tissues.

**Figure 3:**
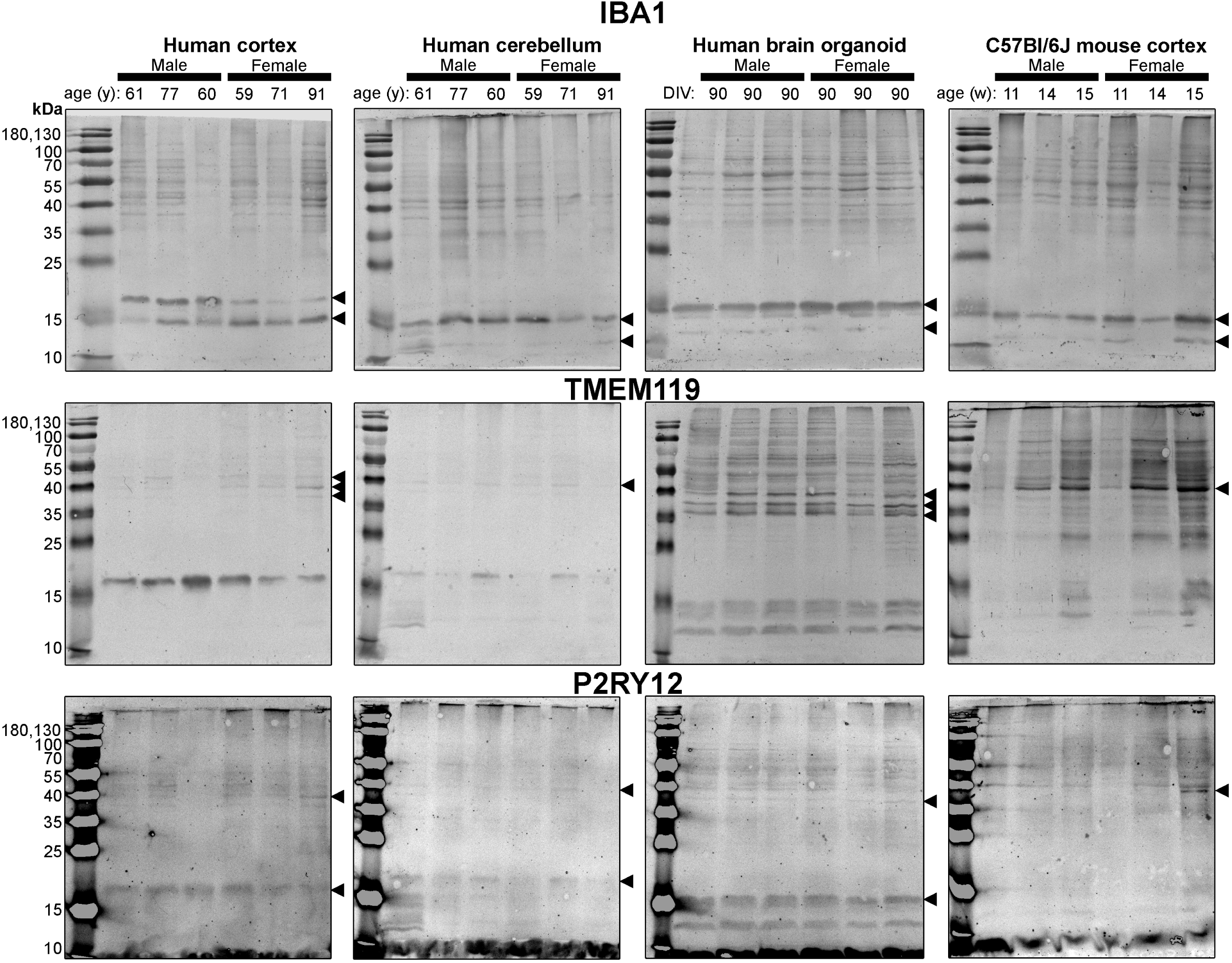
Immunoblots of microglia-associated proteins IBA1 (*top*), TMEM119 (*middle*) and P2RY12 (*bottom*). Samples are the same the protein homogenates described in Figure 2, and the experiment was conducted in the same manner so the immunodensity of bands are visually comparable from one blot to the next. Arrowheads indicate the specific bands used for estimations of protein expression for Figures 6-7. *Abbreviations: days in vitro (DIV), week (w), year (y), ionized calcium-binding adapter molecule 1 (IBA1), transmembrane protein 119 (TMEM119), purinergic receptor P2Y12 (P2RY12)*.

**Figure 4:**
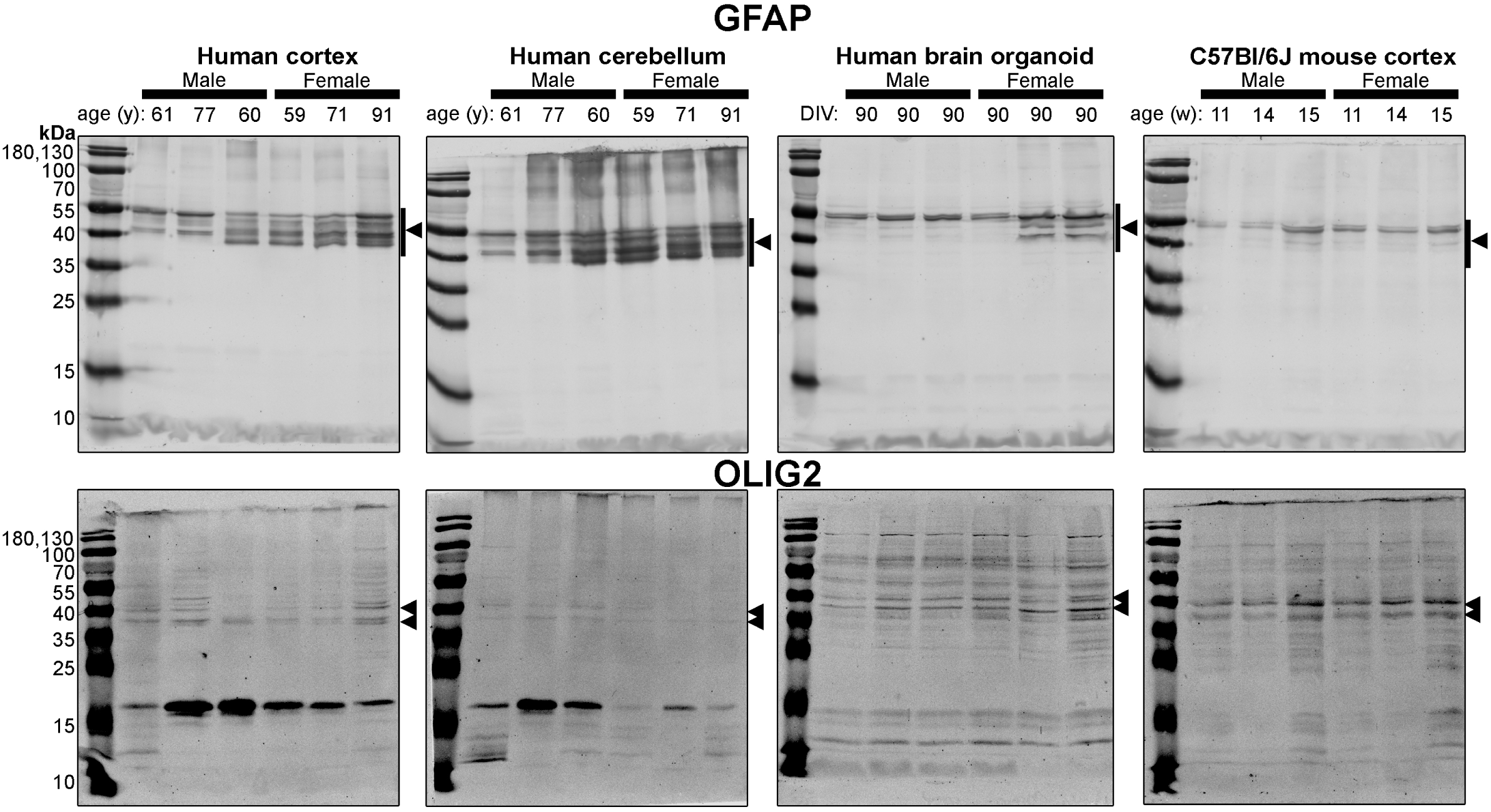
Immunoblots of macroglia (astrocyte and oligodendrocyte)-associated proteins GFAP (*top*) and OLIG2 (*bottom*). Samples are the same the protein homogenates described in Figure 2, and the experiment was conducted in the same manner so the immunodensity of bands are visually comparable from one blot to the next. Arrowheads indicate the specific bands used for estimations of protein expression for Figures 6-7. *Abbreviations: days in vitro (DIV), week (w), year (y), glial fibrillary acidic protein (GFAP), oligodendrocyte transcription factor 2 (OLIG2)*.

**Figure 5:**
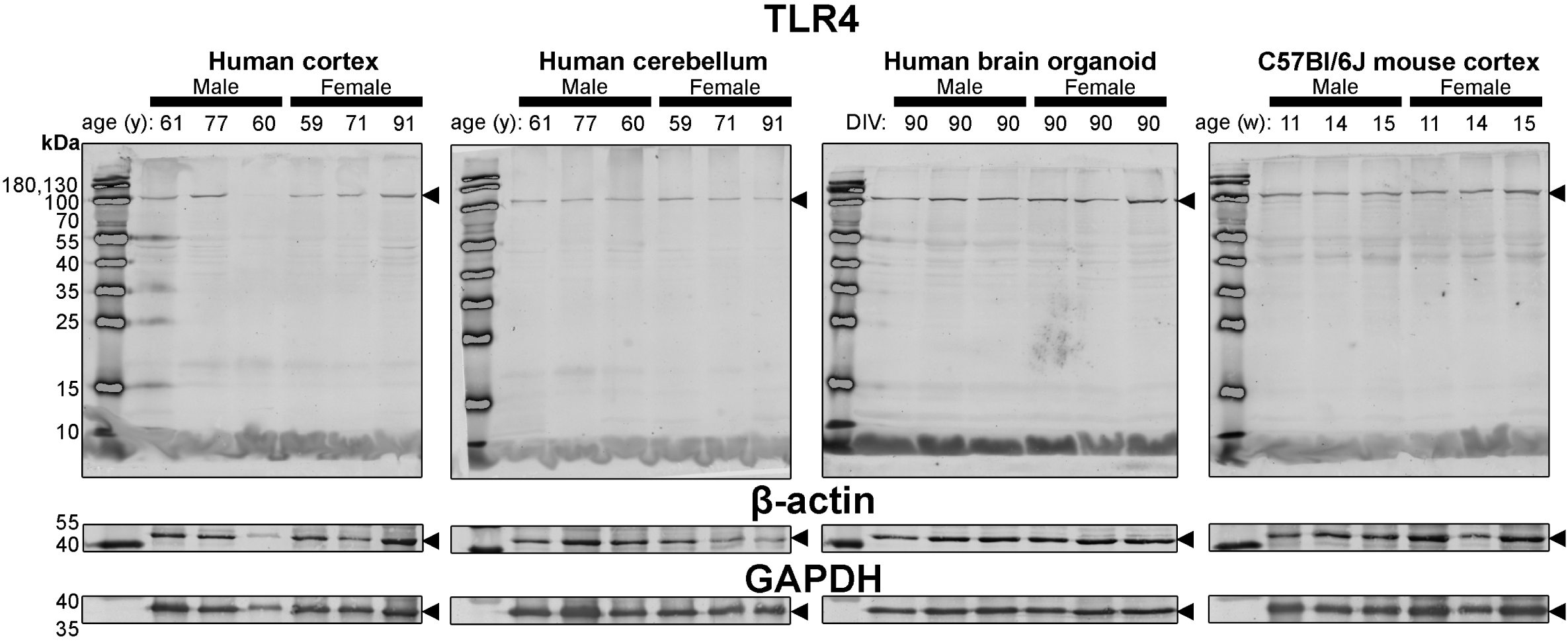
Immunoblots of TLR4 (*top*) as well as housekeeping proteins β-actin (*middle*) and GAPDH (*bottom*). Samples are the same the protein homogenates described in Figure 2, and the experiment was conducted in the same manner so the immunodensity of bands are visually comparable from one blot to the next. β-actin and GAPDH blots are cropped as only one band was detected (see Figure 1). Arrowheads indicate the specific bands used for estimations of protein expression for Figures 6-7. *Abbreviations: days in vitro (DIV), week (w), year (y), toll-like receptor 4 (TLR4), glyceraldehyde-3-phosphate dehydrogenase (GAPDH)*.

Lastly, we probed for TLR4 as well as the housekeeping genes β-actin and GAPDH. TLR4 was included in our list of select proteins as it has been described to have differential levels of expression in mice and human tissue (Smith and Dragunow, 2014). β-actin, TLR4, and TUBB3 exhibit similar protein banding patterns between all tissues tested (**Figure 3**).

### 3.3 Differences and similarities of select protein levels in brain organoids as well as in human and mouse brain parenchymal tissues

Densitometry of immunoblots shown in **Figures 2-5** are summarized in **Figure 6**. When pooling fetal sex, we demonstrate that the protein immunodensity of NeuN (55 kDa), TUBB3 (**Figure 6A**), IBA1, TMEM119, P2RY12 (60 kDa and 25 kDa) (**Figure 6B**), GFAP, OLIG2 (**Figure 6C**), and β-actin (**Figure 6D,ii**) are similar between BOs and human Ctx tissues. The immunodensity of TMEM119 (**Figure 6B,ii**) and TLR4 **(Figure 6D,i**) are more similar between BOs and mouse Ctx tissues, as the human Ctx and Cb homogenates have comparatively less TMEM119 and TLR4. The human Ctx has the highest levels of this 110 kDa NeuN protein compared to other tissue tested (**Figure 6A,i**). Human and mouse Ctx tissues have higher levels of SYN1 than BOs (**Figure 6A,iv**). We also observe that the mouse Ctx has significantly lower levels of GFAP than the human parenchymal tissues but at levels similar to BOs, and OLIG2 immunodensity is lower in the human Cb compared to the human and mouse Ctx (**Figure 6C**).

**Figure 6:**
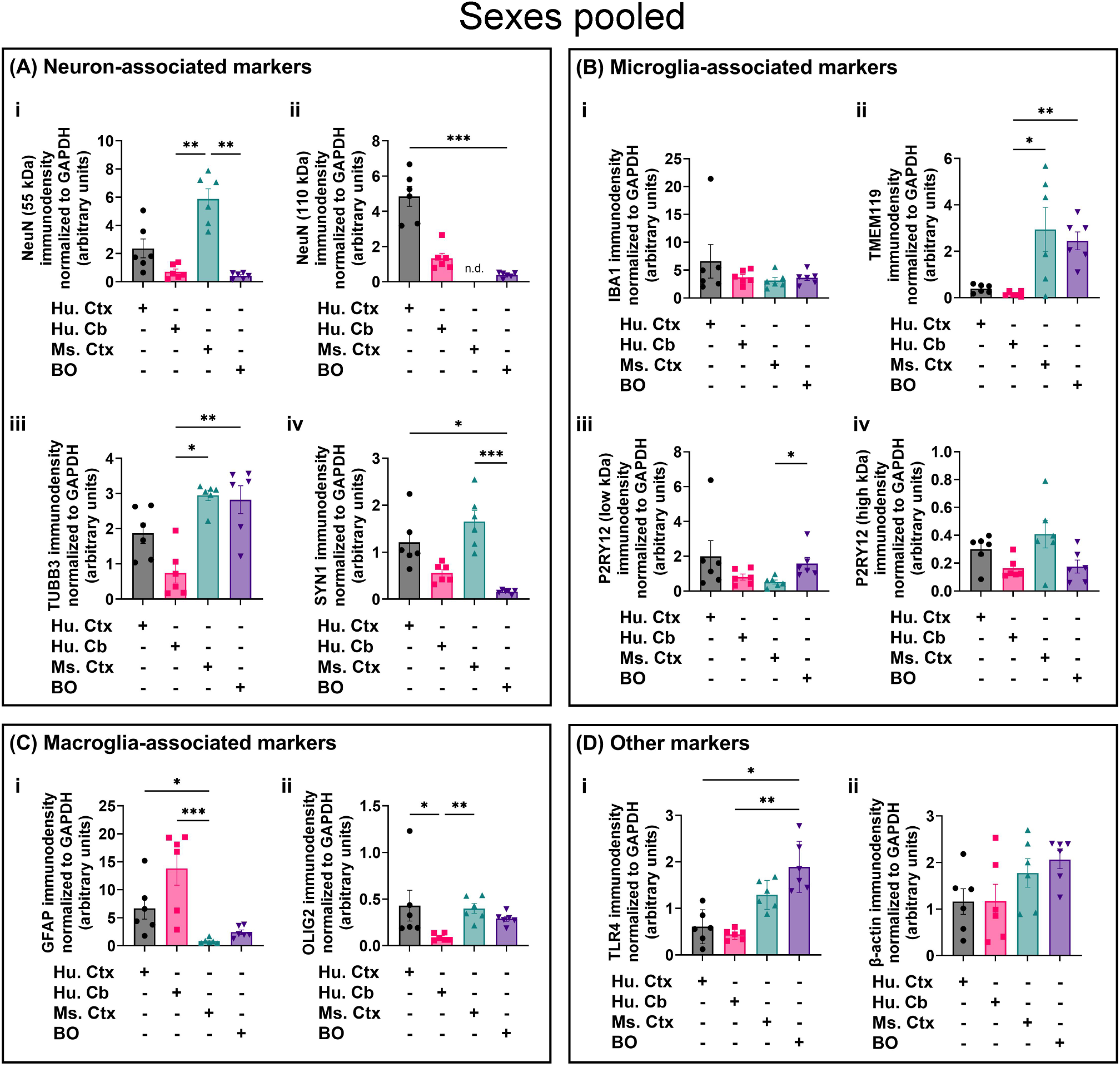
Densitometry of immunoblots (n = 3 males + 3 females, pooled) shown in Figure 3 comparing the expression levels of **(A)** neuron-associated markers (i, NeuN-55 kDa, Kruskal-Wallis: *p = 0.0004*, H = 17.97; ii, NeuN-110 kDa, Kruskal-Wallis: *p < 0.0004*1 H = 15.16; iii, TUBB3, Kruskal-Wallis: *p = 0.0018*, H = 15.03; iv, SYN1, Kruskal-Wallis: *p = 0.0002*, H = 19.21), **(B)** microglia-associated markers (i, IBA1, Kruskal-Wallis: *p = 0.72*, H = 1.31; ii, TMEM119, Kruskal-Wallis: *p = 0.0033*, H = 13.70; iii, P2RY12-25 kDa, Kruskal-Wallis: *p = 0.023*, H = 8.57; iv, P2RY12-60 kDa, Kruskal-Wallis: *p = 0.064*, H = 7.25), **(C)** macroglia (astrocyte and oligodendrocyte)-associated markers (i, GFAP, Kruskal-Wallis: *p = 0.0005*, H = 17.55; ii, OLIG2, Kruskal-Wallis: *p = 0.0023*, H = 14.54), and **(D)** other proteins of interest (i, TLR4, Kruskal-Wallis: *p = 0.0006*, H = 17.46; ii, β-actin, Kruskal-Wallis: *p = 0.13*, H = 5.57). Data presented as means ± SD and normalized to the housekeeping protein GAPDH. Data-points from primary mouse and human tissues were derived from different donors (**Table 2**), while individual BO data-points were derived by pooling five organoids. The five BOs pooled per sample were from the same batch, but individual samples (*i.e.,* data-point) represent a different batch of five organoids (batches defined as BOs generated on different days from iPSCs of a different passage number). * P < 0.05, ** P < 0.01, and *** P < .0001 according to the Dunn’s *post hoc* test.

### 3.4 Sex-dependent differences of select proteins in brain organoids as well as human and mouse parenchymal tissues

Fetal sex has been described to influence the macrostructure of the human parenchyma (DeCasien et al., 2022), yet it is uncertain whether the donor sex of iPSC lines influence the cytoarchitecture of BOs. Thus, we re-analyzed the data shown in **Figure 6** to determine if any sex differences exist in our samples. The 55 kDa NeuN band intensity was significantly less in female mouse Ctx tissue than male mice (**Figure 7A,i**), and GFAP immunodensity was significantly higher in female human Cb tissues than male Cb tissues (**Figure 7C**,**i**). No sex-specific differences were observed for any of the other measure proteins.

**Figure 7:**
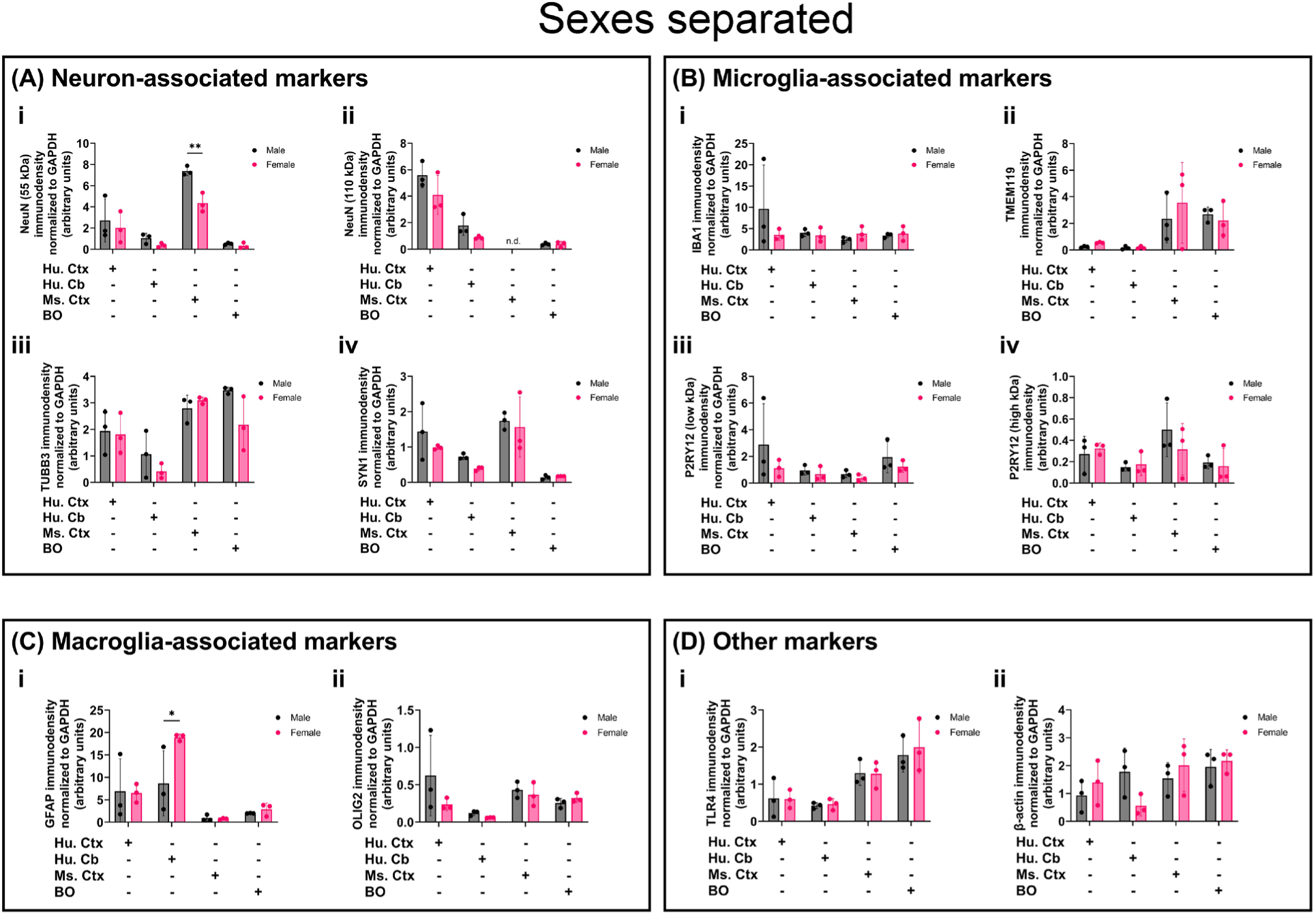
Separation of data-points shown in Figure 2 by fetal sex to compare sex-dependent expression levels of **(A)** neuron-associated markers (i, NeuN 55 kDa; ii, NeuN 110 kDa; iii, TUBB3; iv, SYN1), **(B)** microglia-associated markers (i, IBA1; ii, TMEM119; iii, P2RY12 25 kDa; iv, P2RY12 60 kDa), **(C)** macroglia (astrocyte and oligodendrocyte)-associated markers (i, GFAP; ii, OLIG2), and **(D)** other proteins of interest (i, TLR4; ii, β-actin). Data presented as means ± SD and normalized to the housekeeping protein GAPDH. Data-points from primary mouse and human tissues were derived from different donors (**Table 2**), while BO data-points were derived by pooling five organoids from different batches (batches defined as BOs generated on different days from iPSCs of a different passage number). * P < 0.05 and ** P < 0.01 according to the Sidak’s post hoc test.

### 3.5 Brain organoids display synchronous neural activity at in vitro day 90

We then performed multielectrode array experiments to test whether these BOs exhibited neural activity at *in vitro* day 90 in a pilot experiment. Two weeks after plating the organoids, we detected spontaneous and synchronized network electrical activity (**Figure 8**).

**Figure 8:**
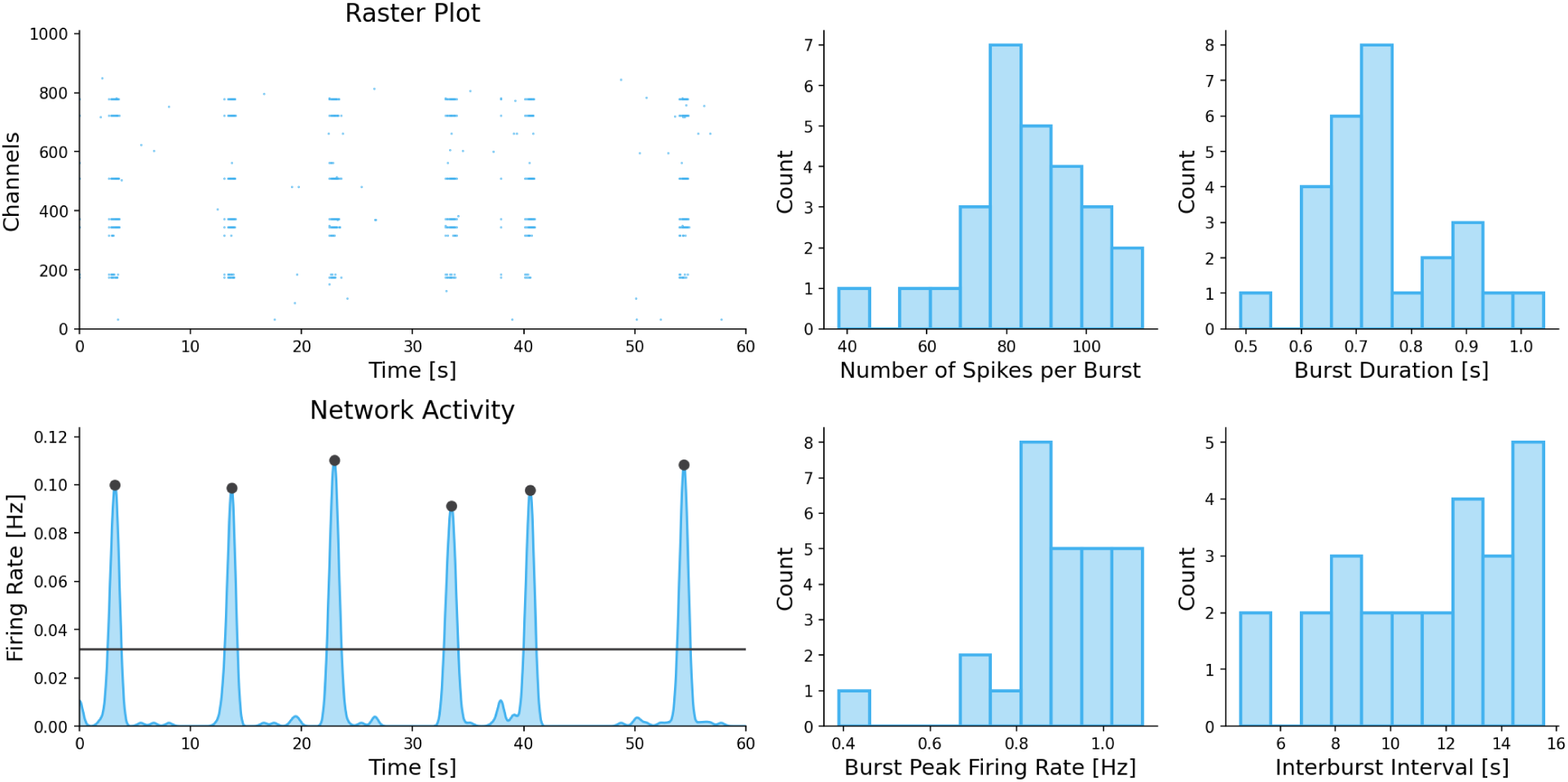
Two weeks after plating, the multielectrode array detects neural network activity in 86i (male) BOs cultured for 90 days *in vitro*. Spontaneous actional potentials (spikes) and network burst electrical activity were observed (*left*), and the electrophysiological properties of this network activity is summarized (*right*).

## 4 Discussion

BOs have been described to be human brain-like tissues that exhibit batch- and cell line-dependent variability (Kim et al., 2021), but studies using stringent protocols demonstrate the variability may not be as pronounced as originally hypothesized given adequate study design (Jourdon et al., 2023; Velasco et al., 2019). A majority of studies have compared the transcriptome of BOs to those of adult and fetal parenchymal tissues (Velasco et al., 2019), but these analyses often do not account for the different proteoforms (e.g., splice isoforms and post-translational modifications) of any given protein. Thus, it is not known whether the transcripts of BOs produce similar proteoforms as the human parenchyma. By extension, it is uncertain whether BOs express proteins in a more human-like or mouse-like manner. Protein-based assays are needed to be adopted in BO studies, as it is well-documented that mRNA levels do not always corroborate levels of the gene product (Arber et al., 2021). A study that demonstrates the protein banding of BOs are more similar to human parenchymal tissues than mouse parenchymal tissues would further validate this platform as a suitable model of the human brain, and support future studies that are interested in investigating sex differences and modulating post-translational mechanisms or splice isoforms. Therefore, we 1) characterized the developmental time course of our BOs and their variability, 2) compared the protein banding patterns of BOs to human Ctx and Cb tissues as well as mouse Ctx tissues, and 3) investigated whether the data from this proof-of-concept study highlighted sex differences.

Our data demonstrate that BOs derived from familial iPSCs (healthy donors) develop proteins at similar time-points, supporting other studies that show cell composition of BOs are predominately driven by the genetic differences of iPSCs from different families (Jourdon et al., 2023). Our data further corroborates the findings of Jourdon et al. (2023) by showing minimal batch-to-batch variability as measured by several different assays, including total protein, resazurin reduction, and expression of housekeeping genes. Importantly, this reproducible consistency occurs without compromising the original visual developmental hallmarks of BOs (Lancaster and Knoblich, 2014). It is essential that studies are able to consistently replicate the cell composition of BOs, as inconsistent composition of cultures will likely bias data interpretation (Wenzel et al., 2023). For example, studies that suggest BO variability is a major hurdle are the same studies that also struggle to generate BOs with consistent protein profiles (Hernández et al., 2022). Stringent standard operating procedures, such as using the same lot number of commercial culture supplements that are known to have batch-to-batch differences, may be essential to reproducibly generating BOs (Chen et al., 2008; Sloan et al., 2018; van der Valk et al., 2010).

Interestingly, the protein profile of BOs exhibits similar, and possibly even less, variability than parenchymal tissues from human donors or clonal mice, as evidenced for example by proteins that are in continual flux such as IBA1. Similarly, BOs also express isoforms of GFAP, and certain human donors displayed similar GFAP banding patterns as the BOs. Consistent with other studies, several isoforms of GFAP were detected in human parenchymal tissues and fewer isoforms were detected in mouse Ctx tissues (Kamphuis et al., 2014, 2012), highlighting interspecies differences. Further studies with a much larger cohort of familial iPSC lines are needed to determine whether BOs have the capacity to retain the donor heterogeneity that we observe in human parenchymal samples, but the transcriptomic study by Jourdon et al. (2023) indicates that at least some donor heterogeneity is retained in BO cultures. It would be exciting if future studies show BOs derived from different donors could capture the heterogeneity of GFAP isoform expression that we observe in human parenchymal tissues. Our data highlight that immunoblotting results of BOs need to be benchmarked against many human brain samples to determine physiological relevance, as there is heterogeneity in the protein banding of parenchymal tissues, which is not often captured by cell lines.

Limited studies have done a comparative immunoblot analysis between human and mouse parenchymal tissues (Bayés et al., 2012; Ishii et al., 2009), complicating the interpretation of immunoblots (Rosell et al., 2020). For example, a NeuN doublet has been reported just above 55 kDa in unguided BOs after 90 days *in vitro* (DIV) and an undisclosed section of adult human and mouse parenchyma (Sriram et al., 2020), which is different than the molecular weight of the doublet we observed in human parenchymal tissues. Similarly, Young et al. (2021) showed TMEM119 antibodies detected a triplet between 38-70 kDa in mouse parenchymal tissues when 15 μg was loaded, and only a 50 kDa band was detected when 3 μg was loaded (Young et al., 2021). In contrast, we observed a triplet with human tissues probed for TMEM119, but only detected a single band in mouse Ctx tissues. It is unclear whether the additional bands at unexpected molecular weights are post-translational modifications, splice variants, or non-specific binding (Rosell et al., 2020). It is also possible that certain bands, such as the 17 kDa bands we observed with OLIG2 and TMEM119 antibodies, are breakdown products that bind together, as demonstrated in other studies (Haytural et al., 2019). Regardless of whether the protein bands detected in our study are proteoforms or non-specific binding, our data supports that BOs are a human-like tissue as they demonstrate protein banding similar to human parenchymal tissue.

While there is well-documented sex differences in regional brain volume, but the cytoarchitectural basis for these differences are unclear (DeCasien et al., 2022). Some reports of sex-dependent abundance of brain cells such as murine microglia (Bordt et al., 2020), while other studies indicate consistent cell proportions in human parenchymal tissue (Johansen et al., 2022). Complicating studies investigating the basis of the sex-dependent brain volume differences is the correlation between cell type proportion and age (Chen et al., 2023). Our data largely supports studies demonstrating a lack of sex-dependent differences in human brain cell proportions (DeCasien et al., 2022; Johansen et al., 2022), and we further show that BOs themselves do not seem to exhibit sex differences in cell composition. Interestingly, we do see a sex difference in the levels of NeuN in mouse Ctx tissue, but no differences in the abundance of murine microglia as Bordt et al. (2020) described in their review. We did, however, observe a sex-dependent difference in GFAP immunodensity in human Cb tissues, but the underlying basis for this difference is unclear. Intriguingly, murine models of traumatic brain injury indicate that GFAP levels are modulated in a sex-dependent manner, indicating that there may have been external factors contributing to our observed sex-dependent differences in GFAP (Sass et al., 2021; Wright et al., 2017). More studies are needed to support whether the sex-dependent differences we observed can be generalized to a wider human population.

Similar to other studies using BOs (Fair et al., 2020; Sharf et al., 2022; Van Lent et al., 2023), we also detect spontaneous and synchronized neural network activity. Studies generating unguided BOs observe synchronized activity between five to six months in culture (Fair et al., 2020; Sharf et al., 2022), while other studies could not detected spontaneous synchronicity without the presence of microglia-like cells (Popova et al., 2021). Since microglia-like cells facilitate the maturation of neural networks in BOs (Popova et al., 2021), it is possible the microglia-like cells in our BOs are underpinning our ability to detect synchronized neural network activity at three months in culture. Interestingly, the electrophysiological parameters of unguided BOs are inconsistent between studies (Fair et al., 2020; Popova et al., 2021; Sharf et al., 2022). Fair et al. (2020) detected spontaneous burst activity every 25 seconds, whereas Sharf et al. (2022) observed activity every five seconds and every 125 seconds in Popova et al. (2021). BOs generated using alternative protocols exhibited a slow firing rate in line with the results by Popova et al. (2021, with burst activity being detected every 80 seconds (Van Lent et al., 2023). The electrophysiology parameters of our BOs were similar to Fair et al. (2020) and Sharf et al. (2022), as we detected burst activity approximately every 12 seconds. The reason underlying the electrophysiological differences between organoid studies is unclear, but it may be due to differences in the cell compositions of the generated BOs. Future studies should consider characterizing the cell compositions of their BOs in each experiment (Wenzel et al., 2023)

We identified that human BOs display protein banding more similar to human Ctx or Cb tissues than mouse Ctx tissues, indicating that human protein proteoforms are conserved in BOs. We further demonstrate that BOs can be generated with minimal batch-to-batch variability in cell composition and metabolism when using iPSCs from healthy male and female siblings. These observations are critical to future studies using BOs, as it demonstrates that immunoblotting results may need to be benchmarked against human parenchymal lysates to be interpreted satisfactorily and it sets a baseline for investigations in sex-dependent responses of cell types to exogenous factors.

## Acknowledgements

TW is supported by funding from the Natural Sciences and Engineering Research Council of Canada through a postdoctoral award. TW and DM further acknowledge financial contributions from the Saskatchewan Health Research Foundation, the Alzheimer Society of Saskatchewan and the University of Saskatchewan. We thank Adam. A. Hussain for their help generating the images shown in Figure 1B during their Honor’s project.

## Author Contributions

**Tyler J. Wenzel:** Conceptualization, Funding Acquisition, Methodology, Investigation, Formal Analysis, Visualization, Writing – Original Draft, Writing – Review & Editing. **Darrell D. Mousseau:** Conceptualization, Funding Acquisition, Writing – Review & Editing.

